# Labile carbon limits late winter microbial activity near Arctic treeline

**DOI:** 10.1101/2020.04.24.058198

**Authors:** Patrick F. Sullivan, Madeline C. Stokes, Cameron K. McMillan, Michael N. Weintraub

## Abstract

It is well established that soil microbial communities remain active during much of the Arctic winter, despite soil temperatures that are often well below −10°C^1^. Overwinter microbial activity has important effects on global carbon (C) budgets^2^, nutrient cycling and vegetation community composition^3^. Microbial respiration is highly temperature sensitive in frozen soils, as liquid water and solute availability decrease rapidly with declining temperature^4^. Thus, temperature is considered the ultimate control on overwinter soil microbial activity in the Arctic. Warmer winter soils are thought to yield greater microbial respiration of available C, greater overwinter CO_2_ efflux and a flush of nutrients that could be available for plant uptake at thaw^3^. Rising air temperature, combined with changes in timing and/or depth of snowpack development, is leading to warmer Arctic winter soils^5^. Using observational and experimental approaches in the field and in the laboratory, we demonstrate that persistently warm winter soils can lead to labile C starvation of the microbial community and reduced respiration rates, despite the high C content of most arctic soils. If Arctic winter soil temperatures continue to rise, microbial C limitation will reduce cold season CO_2_ emissions and alter soil nutrient cycling, if not countered by greater labile C inputs.

---

The importance of overwinter microbial activity to ecosystem C budgets was first recognized in the subalpine forests of Colorado and Wyoming, where relatively deep and often early developing snowpacks maintain soil temperatures of −3 to 0°C throughout the winter^6^. These lightly frozen to unfrozen soils contain substantial amounts of liquid water, as solutes depress the freezing point, minimizing important physical constraints to microbial activity that are present in deeply frozen soils^7^. In these sub-alpine forests, overwinter soil respiration often exceeds 100 g C/m^2^ and is estimated to account for approximately 10% of total annual ecosystem respiration^8,9^. However, sustained microbial activity over the ∼6-month snow-covered season in this relatively warm montane system can lead to late winter labile C limitation of microbial activity^10^. The development of labile C limitation likely serves as an important negative feedback to warming effects on winter ecosystem-atmosphere C flux – limiting the loss of C that would be expected if soil temperature were the primary constraint on microbial activity. Seasonal development of microbial labile C limitation may also have complex effects on overwinter nutrient cycling and availability to plants early in the growing season.

In most arctic ecosystems, the combination of very cold air temperatures and shallow snowpacks leads to soils that are deeply frozen throughout most of the snow-covered season. For instance, at Toolik Lake Field Station on the North Slope of Alaska, soil temperature at 10 cm depth was at or below −5°C for an average of 130 days/year between 1999 and 2017^11^. In these deeply frozen soils, the presence of liquid water is restricted to thin films surrounding soil particles, where the direct and indirect effects of temperature are expected to hold microbial activity at low enough levels to prevent exhaustion of labile C inputs made during the short, but productive arctic growing season^4^. However, the Arctic is warming at twice the rate of the rest of the globe and the greatest changes have been observed during the winter months^12^. Long-term air temperature measurements made in Kotzebue, AK (1943-2019) show a rate of winter warming (+0.7°C /decade, December-February) that is more than twice the rate of summer warming (+0.3°C /decade, June-August, Fig.1). Changes in winter air temperature have been even more dramatic in recent years. Measurements made in our Agashashok River study area (∼65 km north of Kotzebue) show a statistically significant winter warming trend of +4.4°C/decade that corresponds with a non-significant summer warming trend of +0.8°C/decade over the past 14 years. This asymmetric warming of the Arctic may be generating biological responses in winter that outpace corresponding changes in the summer.

**Fig. 1.**
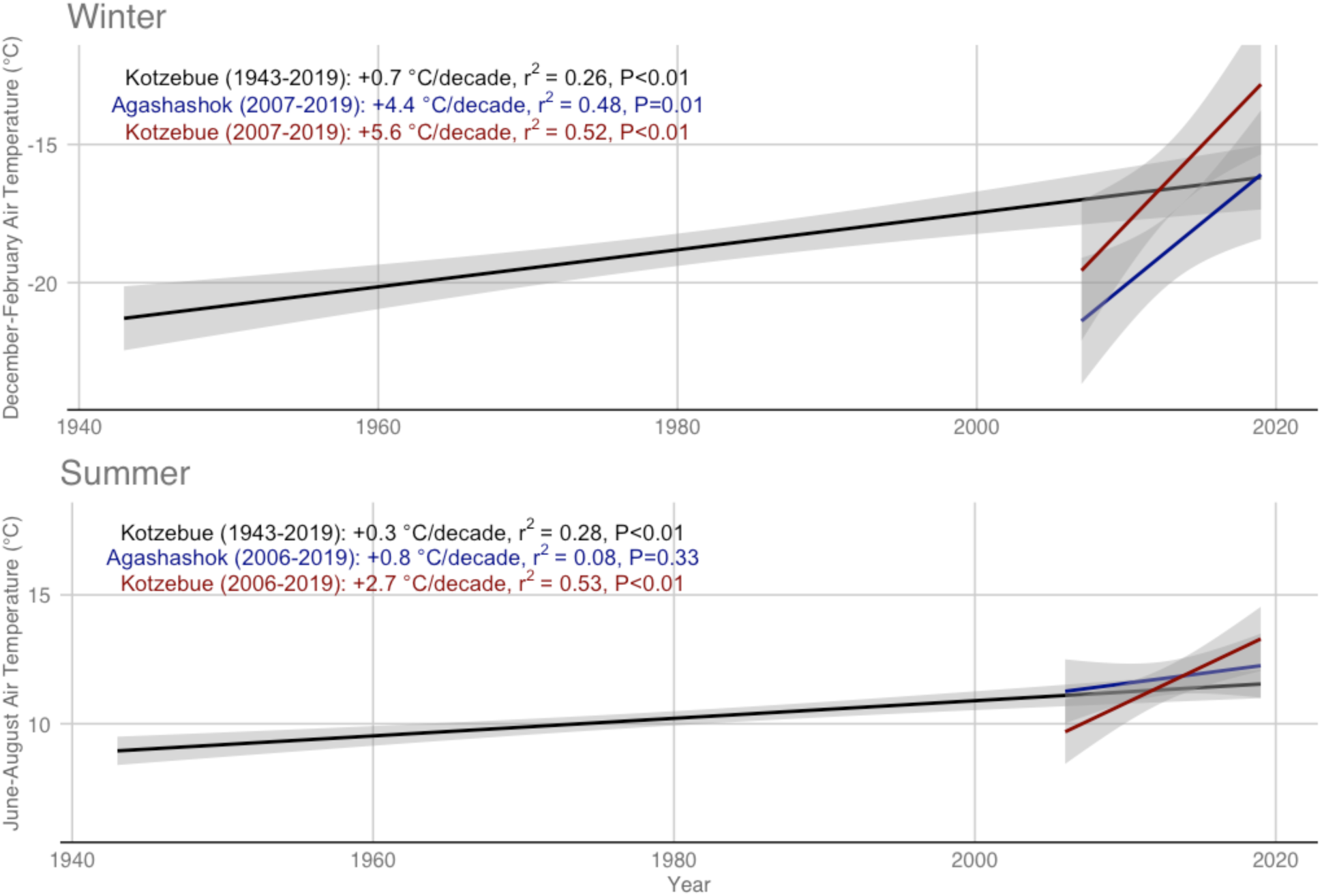
Asymmetric warming in the western Brooks Range of Alaska. Long-term air temperature data from Kotzebue, AK (1943-2019) show more than twice the rate of warming in winter (December-February) than during summer (June-August). More recent data (2006-2019) from our study area in the Agashashok watershed show even greater asymmetry, reflecting a period of rapid winter warming that is also apparent in the recent Kotzebue temperature record (∼65 km south of our study area). Our Agashashok study area has a more continental climate than Kotzebue and, therefore, tends to be slightly warmer during June and July and colder during the winter. The year assigned to each winter includes December of the previous year. Grey shading is the S.E.M.

Efforts to model and/or upscale winter C efflux from arctic ecosystems to the atmosphere recognize the importance of substrate availability for microbial respiration. For instance, in a Pan-Arctic synthesis of winter CO_2_ flux data, it was shown that leaf area index and gross primary production were important predictors of spatial variation in winter C flux from ecosystems to the atmosphere^2^. Accounting for spatial variation in substrate availability represents an important improvement in our ability to upscale and/model winter C losses from the Arctic, but it does not address the potential for temporal development of labile C limitation over the course of warmer winters. Along those lines, a biogeochemical model was recently parameterized to account for labile C depletion within the thin water films around particles in deeply frozen soils^13^. The aim of this revision was to better reflect the observed rapid decrease in microbial respiration with declining temperature in frozen soils. The revised model allows for temporal development of labile C limitation of microbial respiration at a given soil temperature, but it assumes that a small increase in soil temperature would largely alleviate substrate limitation, as the increase in liquid water would make more labile C available for microbial respiration.

Here, we provide evidence from field observations, field experimentation and laboratory incubations that relatively warm winter soils can lead to pervasive labile C limitation of microbial respiration at our study sites near the Arctic treeline in the western Brooks Range. The winters of 2016/2017, 2017/2018 and 2018/2019 differed strongly in terms of air temperature, snowpack development and soil temperature in three treeline ecotones that differ in soil hydrology (hydric, mesic and xeric). During the winter of 2016/2017, cold air temperature in November, coupled with a late-developing snowpack led to deeply frozen soils throughout the winter (Fig. 2). In contrast, warm air temperatures and early developing snowpacks during the winters of 2017/2018 and 2018/2019 led to warmer soils that were consistently only lightly frozen. Measurements of CO_2_ efflux from soils to the atmosphere made using the diffusion gradient method^14^ in late March of each year showed lower fluxes at a given temperature near the end of the warm winters than in March of 2017, near the end of a relatively cold winter (Fig. 3). Although lower than expected in these relatively warm soils, the CO_2_ fluxes measured in March of 2018 and 2019 were similar in magnitude to measurements made in a sub-alpine forest in Colorado^9^ and far greater than measurements made in the Agashashok study area one-decade earlier^16^. This observational evidence of lower than expected microbial activity near the end of the warm winters could reflect depletion of labile C availability after many months of warm soils conducive to substantial microbial activity.

**Fig. 2.**
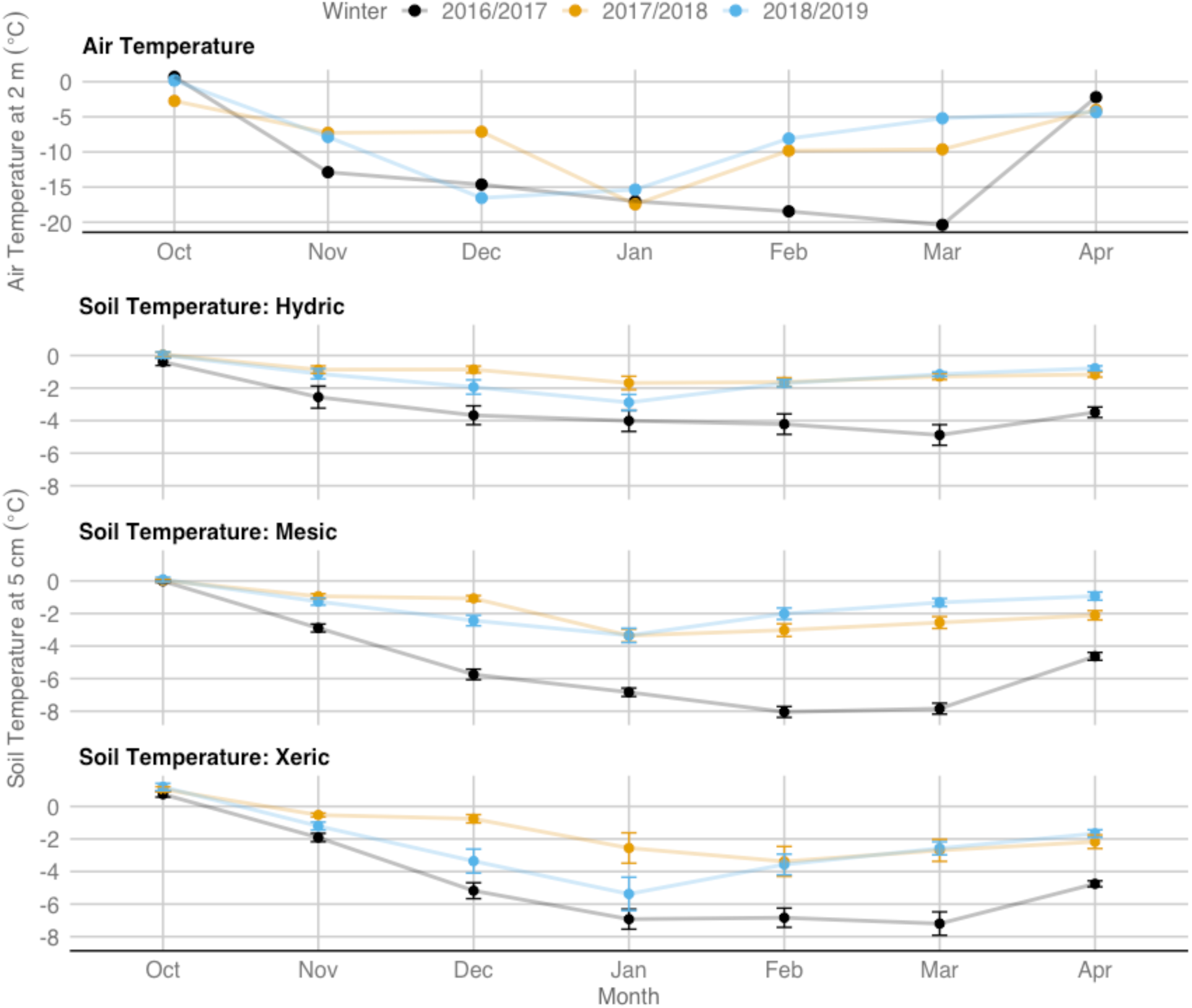
Variation in air and soil temperature across winters. Air temperature was measured at all three sites, but showed negligible variation across sites, as elevation varied by only 35 m. Air temperature data are shown for the Hydric site. Soil temperature is the mean of sensors installed beneath the drip line of 8 control white spruce trees at each site during each winter (Bars= S.E.M.). The air temperature data show much colder conditions during November and February-March in the winter of 2016/2017 than during the unusually warm winters of 2017/2018 and 2018/2019. Soil temperature data show persistently warmer soils during the winters of 2017/2018 and 2018/2019, particularly in February-April.

**Fig. 3.**
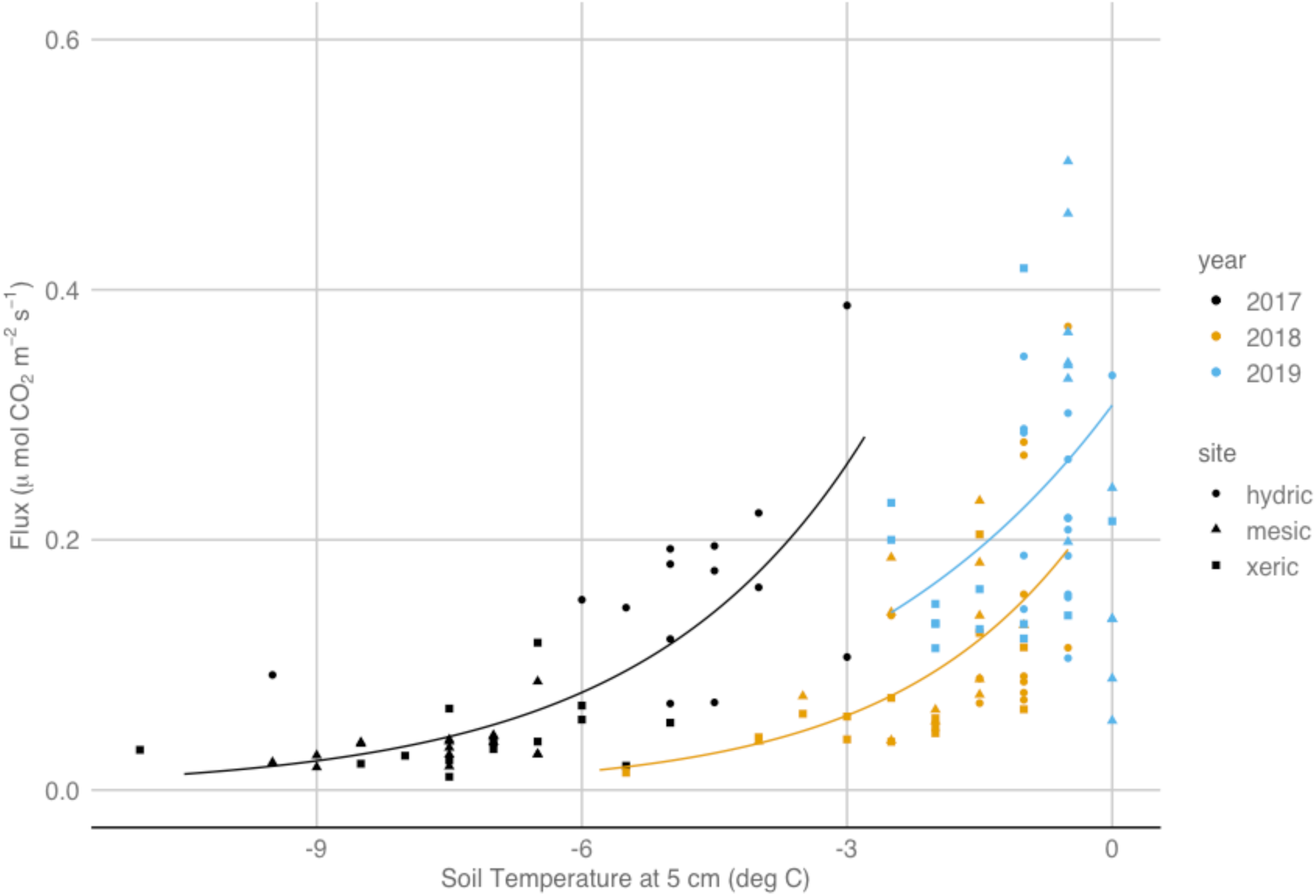
CO_2_ efflux as a function of soil temperature. Measurements were made in three white spruce treeline ecotones in the Agashashok watershed that differ in soil hydrology and soil organic horizon thickness during late March of the contrasting winters of 2016/2017, 2017/2018 and 2018/2019. Q_10_ models were fit separately to data from each year. Measurements were made beneath the dripline of both unmanipulated control trees (n=8/site) and those treated with 1.5 m tall snowfences to increase snow depth (n=8/site). The results show that CO_2_ efflux at a given temperature was greater near the end of the relatively cold winter of 2016/2017.

To further test for labile C limitation of microbial activity, we conducted experiments in the field and in the laboratory. In the field, we carried out a labile C addition experiment, in which we added 100 g C/m^2^ of powdered glucose to the soil surface of five treatment plots, which were paired with nearby control plots, at each of our three treeline sites. Soil surface temperatures were consistent and relatively warm (−2 to −0.5°C) during the field glucose additions. We made pre-treatment measurements of CO_2_ flux from both control and treatment plots and then carefully (and completely) removed the snowpack from both treatment and control plots. After amending the treatment plots with powdered glucose, we immediately returned the snowpack to both control and treatment plots. We tested the optimal timing of treatment response measurements in a boreal forest in Anchorage, Alaska, prior to implementation in the Brooks Range (Fig. S1). Testing revealed a large and still increasing CO_2_ flux response to glucose addition 72 hours after application. Measurements of CO_2_ flux made 72 hours after glucose application at our Brooks Range treeline sites revealed a doubling of CO_2_ flux from an overall control mean of 0.40 to a glucose treatment mean of 0.89 μmol m^-2^ s^-1^ (Fig. 4, F_1,24_=52.5, P<0.01). There was no statistical evidence that the CO_2_ flux response to glucose addition varied across our treeline sites (F_2,22_=0.25, P=0.78), although there was a trend toward a proportionally greater response at the xeric site, which has a much smaller pool of soil organic C than the mesic and hydric sites.

**Fig. 4.**
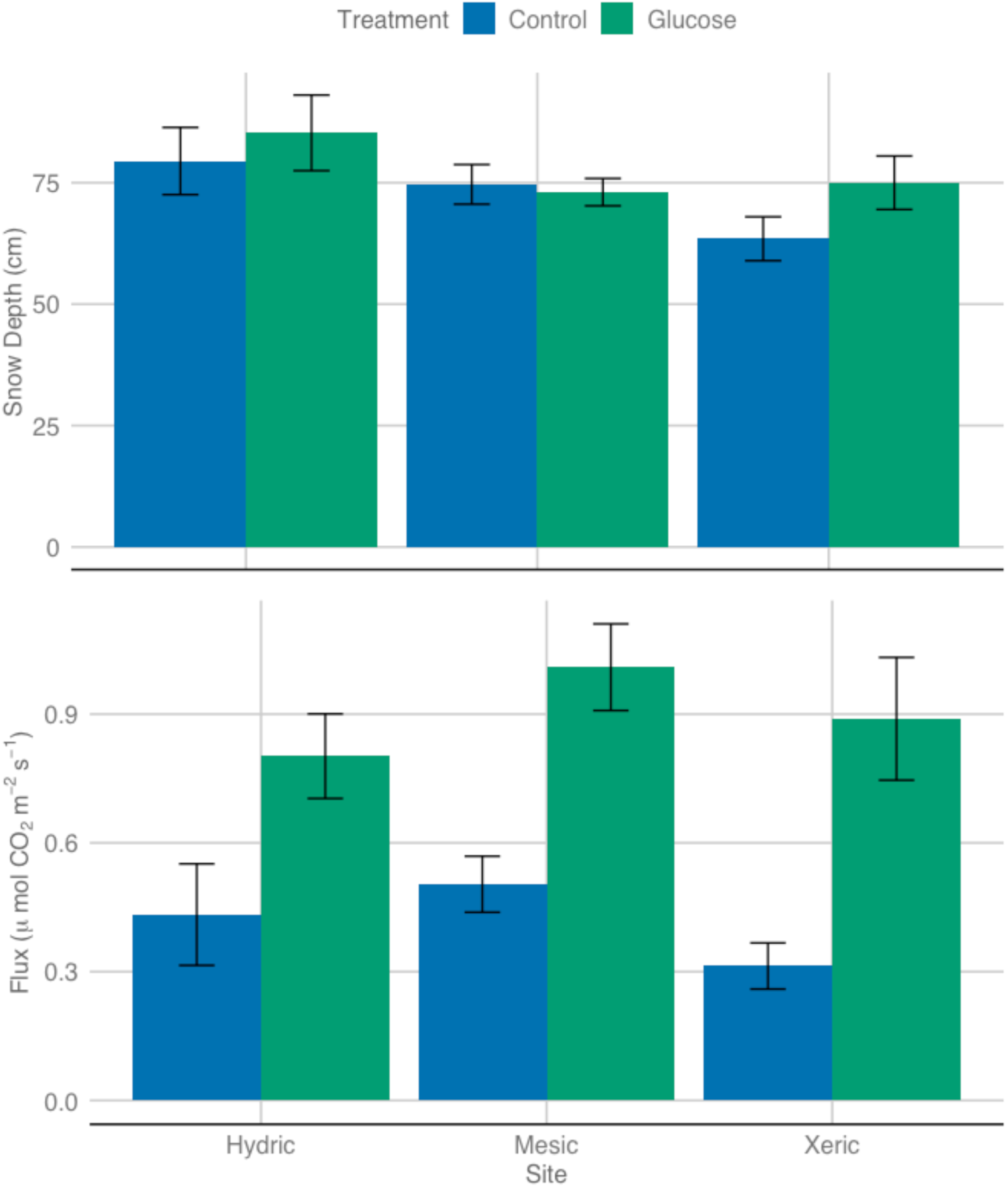
Effects of experimental glucose addition on late winter CO_2_ efflux. Measurements were made 72 hours after applying 100 g C/m^2^ in the form of glucose powder to the soil surface of 5 treatment plots, which were paired with 5 corresponding control plots, at each of three treeline sites in late March of 2019. Soil surface temperatures were consistent and relatively warm (−2 to −0.5°C) during the field glucose additions. Pre-treatment snow depths are shown in the upper panel. Results show strong stimulation of CO_2_ efflux, which is a proxy for microbial activity, with limited evidence of variation in the proportional response across sites. Bars are S.E.M.

In the laboratory, we conducted temperature-controlled incubations of soils from our hydric and xeric sites. Soil temperatures were held at −10, −6, −2, 2 and 6°C and crossed with labile C (cellobiose) additions of 0, 0.2, 0.4 and 2 mg C/g dry soil. Over the three-month incubation, we found strong responses of microbial respiration to labile C addition that depended upon incubation temperature (Fig. 5). Microbial respiratory responses to labile C addition were minimal at −10°C and increased progressively with rising soil temperature, even when the analysis was restricted to subzero soil temperatures (Fig. S2). Respiration, and the effect of C addition on respiration, declined during the three-month incubation, especially at warmer temperatures, indicating rapid utilization of the labile C (Fig. S3).

**Fig. 5.**
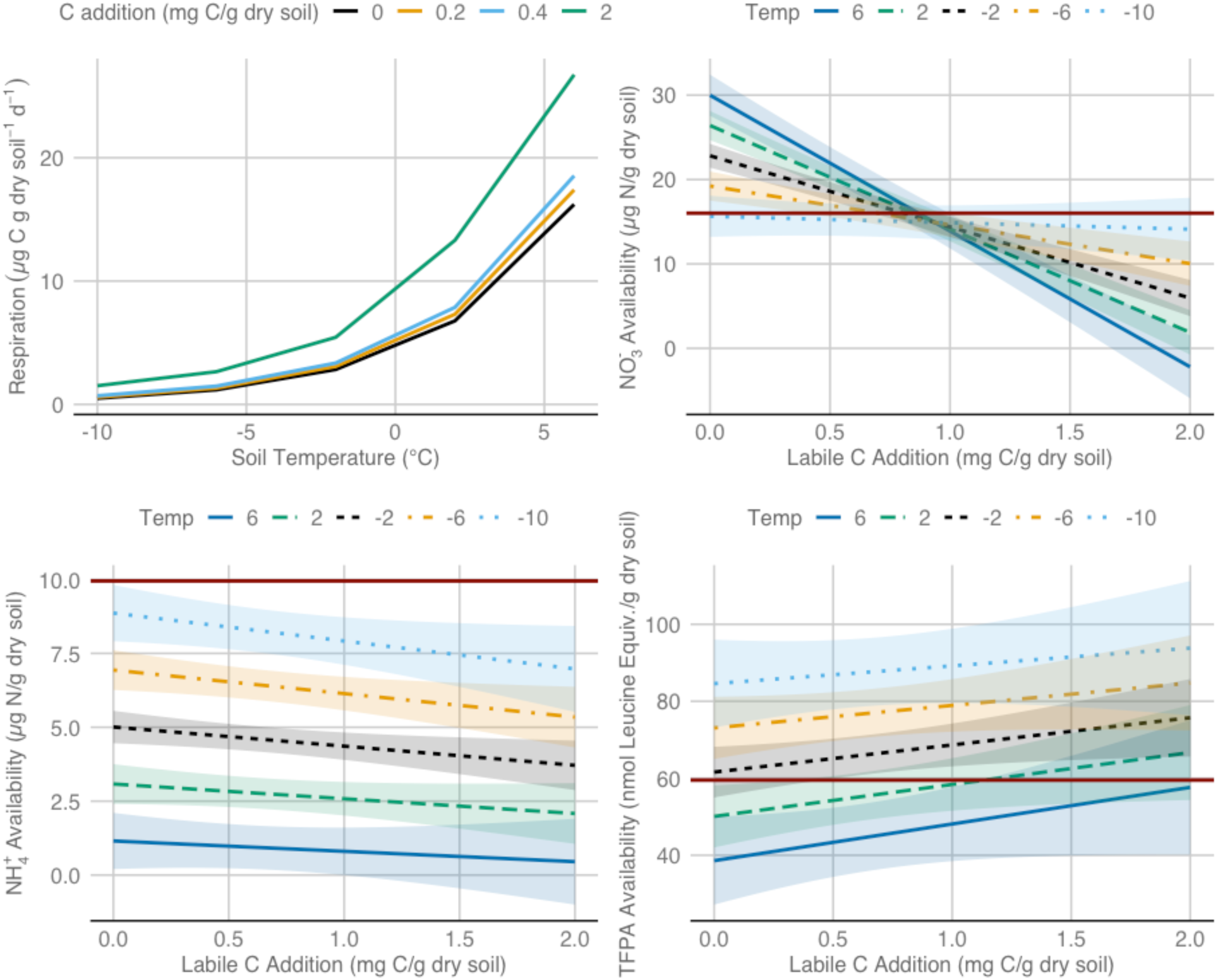
Microbial respiration and soil nutrient availability responses to labile C addition in temperature-controlled laboratory incubations. Cellobiose was added at rates of 0, 0.2, 0.4, and 2 mg C per g dry soil to a homogenized composite of root-free organic soil from our hydric and xeric sites and each was separately incubated at −10, −6, −2, 2, and 6 °C (n=4/temperature*C addition treatment). Respiration was measured at approximately weekly intervals throughout the incubations. Soil NO_3_^-^, NH_4_^+^, and total free primary amine (TFPA) availability was measured at the beginning and end of the three-month incubations. The respiration panel shows predictions of a non-linear mixed effects model that was fitted to the data, while the nutrient availability panels are effects plots from linear models that include temperature, labile C addition and their interaction as predictors. Shading indicates the S.E.M.

Measurements of soil nutrient availability made at the beginning and end of the incubation period suggest that development of labile C limitation will have important implications for overwinter soil nitrogen (N) cycling (Fig. 5, Figs. S5-S8). We also made measurements of orthophosphate-P availability (Fig. S9), which was low overall (near detection limits) and there was no evidence of effects of temperature (t=-1.5, P=0.14), nor labile C availability (t=-1.0, P=0.32). Among the forms of N, NO_3_^-^ availability increased with warming at lower levels of added C (<∼1 mg C/g dry soil), indicating increasing nitrification as a function of temperature, which is consistent with C limitation. NO_3_^-^ declined strongly with labile C addition and the magnitude of decline increased with soil temperature (Temperature*C addition: t=-7.9, P<0.01), indicating that warming in the presence of available C promoted NO_3_^-^ immobilization. Likewise, NH_4_^+^ availability declined with labile C addition (t=-2.0, P=0.05) and with soil temperature (t=-9.8, P<0.01), indicating that increased C and warmer temperatures stimulated immobilization. These results suggest that inorganic N, and especially NO_3_^-^ availability, might increase in response to the development of labile C limitation. Finally, total free primary amine (TFPA) availability, which is a proxy for organic N availability, increased with labile C addition (t=2.2, P=0.03) and decreased with soil temperature (t=-4.9, P<0.01), suggesting that changes in organic N availability may be opposite in direction to those for inorganic N in response to labile C limitation. Collectively, our soil N results suggest that development of labile C limitation will likely have important implications for overwinter N cycling in Arctic soils and may shift the balance of available N forms between NO_3_^-^ and NH_4_^+^ and between inorganic and organic, which will have important implications for microbial stoichiometry and community composition, soil N trace gas emissions and nitrate runoff, plant productivity and vegetation community composition.

Our field observations, field experiments and laboratory incubations consistently indicate that labile C limitation of microbial respiration can develop over time in lightly frozen arctic soils. This discovery has important implications for current estimates and projections of future Pan Arctic C budgets with warming, while adding complexity and uncertainty to relationships among vegetation, snow and soil nutrient cycling at high latitudes. Recent estimates and projections of Pan Arctic winter CO_2_ efflux assume that the temperature response of microbial respiration is static over time^2^. For instance, a soil temperature of −3°C is expected to yield the same CO_2_ flux in March as in November. However, our results suggest that a soil temperature of −3°C in November might yield a higher CO_2_ flux than in March if soils remain relatively warm during the intervening months. Improved modeling of overwinter C efflux will likely require temperature response curves that vary over time with changes in substrate availability. Over longer time periods, labile C limitation of winter microbial respiration could act as an important negative feedback to warming-induced changes in Arctic C budgets. Labile C limitation of microbial respiration may become more common if asymmetric warming leads to an imbalance whereby overwinter increases in microbial activity are not balanced by increases in vegetation productivity and associated labile C production. However, if development of labile C limitation leads to changes in nutrient cycling that yield greater N availability to plants, it is possible that increased vegetation productivity could lead to greater labile C inputs that might prevent further development of labile C limitation.

One of the most widespread recent changes in arctic vegetation is the expansion of tall shrubs into low-statured arctic tundra^17^. In addition to direct effects of growing season climate warming on shrub growth and reproduction, it is thought that, once established, tall shrubs trap snow, which insulates soils in winter and allows for greater overwinter microbial activity, releasing nutrients that are available to support further shrub growth and expansion^3^. This positive feedback loop among tall shrubs, snow and soil microbes is expected to reinforce the process of tall shrub encroachment. A similar process may operate in treeline environments, where the abrupt change in surface roughness between forest and tundra leads to a reduction in wind speed and both reduced sublimation and increased snow deposition among the trees^16^. Interactions among treeline trees, snow and soil microbes thus have the potential to facilitate treeline advance. Our finding that labile C limitation of microbial respiration can develop over time in lightly frozen arctic soils adds complexity and uncertainty to our understanding of relationships among vegetation, snow, microbial activity and soil nutrient cycling in arctic ecosystems. Our discovery of labile C limitation of overwinter microbial activity has the potential to alter understanding of soil temperature feedbacks and interactions and therefore represents an important new avenue for future research in the Arctic and in other ecosystems with seasons of limited photosynthetic labile C production.

## Methods

### Site Description

Field measurements and sample collections were made in three diffuse treeline ecotones near the Agashashok River in the Baird Mountains of northwest Alaska. The study area is near the northern and western limits of white spruce (*Picea glauca*) in Alaska. The northernmost tree in the Agashashok watershed is approximately 12 km north of the study area. The hydric treeline (67.47 N, 162.20 W, 155 m asl) has an understory of wet sedge tundra that is dominated by *Carex bigelowii* and mosses of the genera *Sphagnum* and *Hylocomium* with occasional tussocks of *Eriophorum vaginatum*. The mesic treeline (67.48 N, 162.22 W, 135 m asl) is typical tussock tundra, dominated by *Eriophorum vaginatum, Betula nana, Rhododendron groenlandicum, Vaccinium uliginosum, Salix pulchra* and mosses of the genera *Hylocomium, Pleurozium*, and *Sphagnum*. The xeric treeline (67.47 N, 162.21 W, 170 m asl) has an understory of dry heath tundra. The most common plant species at the xeric site is *Dryas octopetala*, with abundant lichens and lesser amounts of *Cassiope tetragona* and *Empetrum nigrum*. The area is ∼65 km north of Kotzebue, Alaska and is accessed using bush planes in the summer and snowmachines in the winter.

The study sites are part of a snowfence experiment, designed to examine the role of winter snow depth as a driver of treeline tree growth and reproduction. Snowfences (1.5 m tall, 7 m long) were installed 2 m upwind of eight white spruce trees (∼5 cm dbh, ∼3.5 m tall) at each site in early September of 2016. Each of the three sites is equipped with a meteorological station that monitors air temperature and snow depth, along with soil temperature and soil moisture at 10 cm depth increments. iButton temperature loggers (DS1921G) are installed at 5 cm depth beneath the dripline of 8 control and 8 snowfence trees at each site (48 trees total).

Long-term air temperature data for Kotzebue, Alaska were acquired from the National Center for Environmental Information at the National Oceanic and Atmospheric Administration (https://www.ncdc.noaa.gov/). Measurements of air temperature at 2 m height have been made since June of 2006 on a riverside terrace in the Agashashok watershed^18^. Measurements are made using a CS215 sensor (Campbell Scientific, Logan, UT, USA) that is housed within a 6-plate radiation shield (R.M. Young, Traverse City, MI, USA). The sensor is scanned every 15 minutes and hourly averages are logged to a CR1000 datalogger (Campbell Scientific, Logan, UT, USA).

### Field CO_2_ Flux Measurements

The study sites were visited during the final week of March in 2017, 2018 and 2019 and measurements of CO_2_ efflux were made at the dripline of each study tree using the diffusion gradient method^14,16^ during periods with light winds. Measurements of atmospheric [CO_2_] at the snow surface and subnivean [CO_2_] at the ground surface were made at each study tree using a hollow stainless-steel probe that was etched with snow depth increments (Snowmetrics, Fort Collins, CO) and plumbed with mm I.D. polyethylene tubing. The tubing was fitted to the inlet of a LI-840 NDIR CO_2_ and H_2_O analyzer (LI-COR Environmental, Lincoln, NE), which was equipped with a micro-diaphragm pump (850 ml/min, KNF Neuberger Inc., Trenton, NJ) downstream of the optical bench. In addition to measurements of atmospheric and subnivean [CO_2_] at each study tree, measurements were also regularly made of [CO_2_] at 10 cm intervals throughout the snowpack to test for potential barriers to diffusion that could lead to overestimates of CO_2_ efflux when using the 2-point diffusion method^14^.

After completing the [CO_2_] and snow depth measurements at each tree, three snow pits were excavated in representative areas within the treeline ecotone at each site. Measurements of snow density and temperature were made in continuous 10 cm intervals along the walls of the snow pits using a RIP 1 density cutter (Snowmetrics, Fort Collins, CO) and a 600 g capacity spring scale with 5 g resolution (Pensola, Schindellegi, Switzerland). Corresponding measurements of snow temperature were made using a dial stem thermometer.

Diffusion of CO_2_ from the subnivean to the atmosphere was estimated as follows^19^:

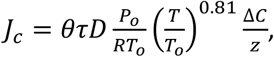

where *J*_*c*_ is CO_2_ efflux (µmol m^-2^ sec^-1^), θ is snowpack porosity (unitless), τ is snowpack tortuosity (unitless), *D* is the diffusion coefficient for CO_2_ in air (0.1381 x 10^−4^ m^-2^ sec^-1^), P_0_/RT_0_ is the molecular density of CO_2_ at standard temperature and pressure (44.613 mol m^-3^), *T* is snowpack temperature (K), Δ*C* is the difference in [CO_2_] between the subnivean and the atmosphere (µmol/mol) and *z* is snow depth (m). Snowpack porosity (θ) was estimated using mean snowpack density (ρ):

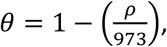

where 973 g/l is the density of ice. Snowpack tortuosity (τ) was also estimated as a function of density^20^:

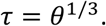

### Field Labile C Addition Experiment

During the final week of March in 2019, 250 g/m^2^ of glucose powder (Product # G8270, Sigma Aldrich, Saint Louis, MS) was added to the soil surface of five 1.0 m^2^ treatment plots, which were paired with five control plots, in each treeline ecotone. Prior to glucose application, the snowpack was carefully removed from a 2.25 m^2^ area centered on each control and treatment plot. After removing the more consolidated upper portion of the snowpack, the large, faceted depth hoar crystals were swept off the plot using a whisk broom. Glucose powder was evenly spread over the central 1.0 m^2^ area of the treatment plots. The snowpack was then returned to the control and treatment plots, first by sweeping the depth hoar back on to the plot and then by returning the blocks of consolidated upper snowpack. The snowpack was lightly packed using snowshoes and any cracks visible at the snow surface were filled with fresh snow. After 72 hours, measurements of CO_2_ efflux were made using the aforementioned diffusion method near the center of each control and glucose treatment plot. After completing the [CO_2_] measurements, snow pits were excavated for measurements of snow density and temperature as described above at each control and treatment plot in each treeline ecotone.

### Temperature Controlled Laboratory Incubations

Organic horizon tundra soils were collected to a depth of 15 cm in summer of 2018 from the hydric and xeric treeline sites. Soils were frozen and shipped to Toledo, OH USA, where they remained frozen until initiation of the incubation experiment. Soils were thawed and homogenized into a single composite, while removing rocks, large roots and woody debris in roughly three parts hydric to one part xeric, reflecting the quantity of available material. Soils were then kept in a 4°C incubator at a near constant soil moisture of ∼50% water holding capacity for roughly three weeks.

To determine the effects of labile C availability and temperature on respiration, the composite soil was incubated with different amounts of labile C at a range of temperatures. Incubations were initiated by placing 25 g (wet mass) of composite soils into 80 half pint wide mouth canning jars (Jarden Corporation, Muncie, IN, USA) fitted with septa, and maintained at 4°C. 5 mL of cellobiose solution at 4°C was added at concentrations of 0, 0.5, 1, or 5 mg C mL^-1^ in reagent grade water (n=20/cellobiose concentration). This approach achieved C additions of 0, 0.2, 0.4, and 2 mg C per g dry soil, respectively. Samples of each C addition treatment were loosely covered with jar lids to prevent CO_2_ buildup and immediately placed in lab incubators set at −10, −6, −2, 2 and 6°C (n=4/temperature*C addition).

A LI-820 infrared gas analyzer (LI-COR Biosciences, Lincoln, NE, USA) outfitted for static injections was used to measure respiration every 3-4 days in the first month, once a week in the second month, and every other week thereafter. To measure respiration, samples were uncovered and vented using a desk fan to clear the headspace and then sealed. To allow enough time for detectable concentrations of CO_2_ to accumulate in jar headspaces, the samples at below-freezing temperatures had to be incubated for 3-4 days, while those at above-freezing temperatures only required 3 hours. While measuring respiration from below freezing incubations, jars were kept on ice. A minimum of two 2 ml headspace samples per jar were analyzed (a third was analyzed if the first two were inconsistent). CO_2_ concentrations were estimated using a calibration curve created with 2500 and 5000 ppm standards, and respiration is reported as μg C g dry soil^-1^ day^-1^.

At the end of the incubation, soils were extracted for carbon and nutrient analysis. 5 g of soil was combined with 25 mL of 0.5 M K_2_SO_4_ in a 50 mL tube and placed on an orbital shaker for 1 hour. The sample was then filtered using a Whatman #1 paper filter (2 µm pore size) and a vacuum filtration system^21^. Total reducing sugar (TRS) concentrations (Fig. S4) in the unfumigated extracts were determined in four analytical replicates per sample using a colorimetric microplate assay with glucose as the standard and expressed in terms of glucose equivalents^22,23^. NH_4_^+^-N and NO_3_^-^-N were measured on three analytical replicates per sample with colorimetric microplate assays^24,25^. Total free primary amines (TFPA) were quantified using a fluorometric microplate assay with leucine as the standard and expressed in terms of leucine equivalents^26,27^.

Microbial biomass (Figs. S10 and S11) was determined using a modification of the chloroform fumigation extraction method^28,29^, in which 2 mL of chloroform was added to 5 g of soil in a 250 mL Erlenmeyer flask, and then immediately stoppered. After 24 hours the stoppers were removed, and the chloroform vented in a fume hood for 30 minutes. Samples were then extracted as described above.

Dissolved organic C (DOC) and total dissolved nitrogen (TDN) concentrations were measured using a Shimadzu total organic C (TOC-V_CPN_) analyzer with a total nitrogen (TN) module (Shimadzu Scientific Instruments Inc., Columbia, MD, USA). Extractable orthophosphate-P was measured on three analytical replicates per sample with a colorimetric microplate assay^30^. All microplate assays were read on Bio-Tek Synergy HT microplate reader (Bio-Tek Inc., Winooski, VT, USA). TOC, TN, and P concentrations in unfumigated extracts were subtracted from those in fumigated extracts to estimate extractable microbial biomass C, N, and P. Extractable microbial biomass C, N, and P were not corrected for extraction efficiency, which is unknown in these soils, and are expressed as μg (C, N, or P) g^-1^ dry soil.

### Statistical Analyses

All data processing, statistical analyses and data visualization were performed using R 3.6.1^31^. Q_10_ models with the following form were fit to tree-level soil temperature and CO_2_ efflux estimates across sites, separately for each year, using the nls function in R:

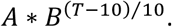

The effect of experimental glucose addition to field plots was examined using a linear model with the pre-treatment CO_2_ flux, site (hydric, mesic, xeric) and treatment (control, glucose) as the main effects. Interactions between site and treatment (F_2,22_=0.25, P=0.78) and between pre-treatment CO_2_ flux and treatment (F_1,23_=0.30, P=0.59) were tested and excluded from the final model.

Respiration data from the temperature-controlled laboratory incubations were analyzed using a nonlinear mixed effects model in the nlme package^32^. Model parameters were included for the level of labile C addition, a Q_10_ temperature response function (as described above) and the interaction between labile C addition and the Q_10_ temperature response function. All three parameters were allowed to vary with sample, which was repeatedly measured over time during the incubation. The predict function in R was used to make predictions of respiration at each level of labile C addition, for each temperature. To examine changes in respiration and changes in the effect of C addition on respiration over time in the incubation, we split the data approximately into two halves: early and late. The nonlinear mixed effects model was then fit separately to the early and late data and respiration predictions were made as described above. In order to fit the model to the data from late in the incubation, we were forced to simplify the random effects, such that only *A* in the Q_10_ model was allowed to vary across samples. There was limited evidence of an interaction between temperature and labile C addition in the late incubation data, but we retained the interaction in the model to improve comparability of predictions with the early incubation data.

Measurements of soil nutrient availability from the end of the incubations were analyzed using linear models including temperature, labile C addition level and their interaction. Interaction plots were generated using the interact_plot function in the interactions package^33^. All graphics were produced using the ggplot2 package^34^.

### Data Availability

The datasets generated during this study will be submitted to the Arctic Data Center (https://arcticdata.io/) upon acceptance of the manuscript for publication. In the meantime, they are available from the corresponding author upon request.

## Acknowledgements

This study was funded by United States National Science Foundation awards OPP-1504538 to P.F.S. and OPP-1503939 to M.N.W. Golden Eagle Outfitters provided valuable logistical support. A. Brownlee assisted with fieldwork.

## Author Contributions

P.F.S. and M.C.S. conducted the fieldwork. C.K.M. performed the laboratory incubations under the supervision of M.N.W. P.F.S. analyzed the data and drafted the manuscript. All authors contributed to revisions.

## Extended Data

**Fig. S1.**
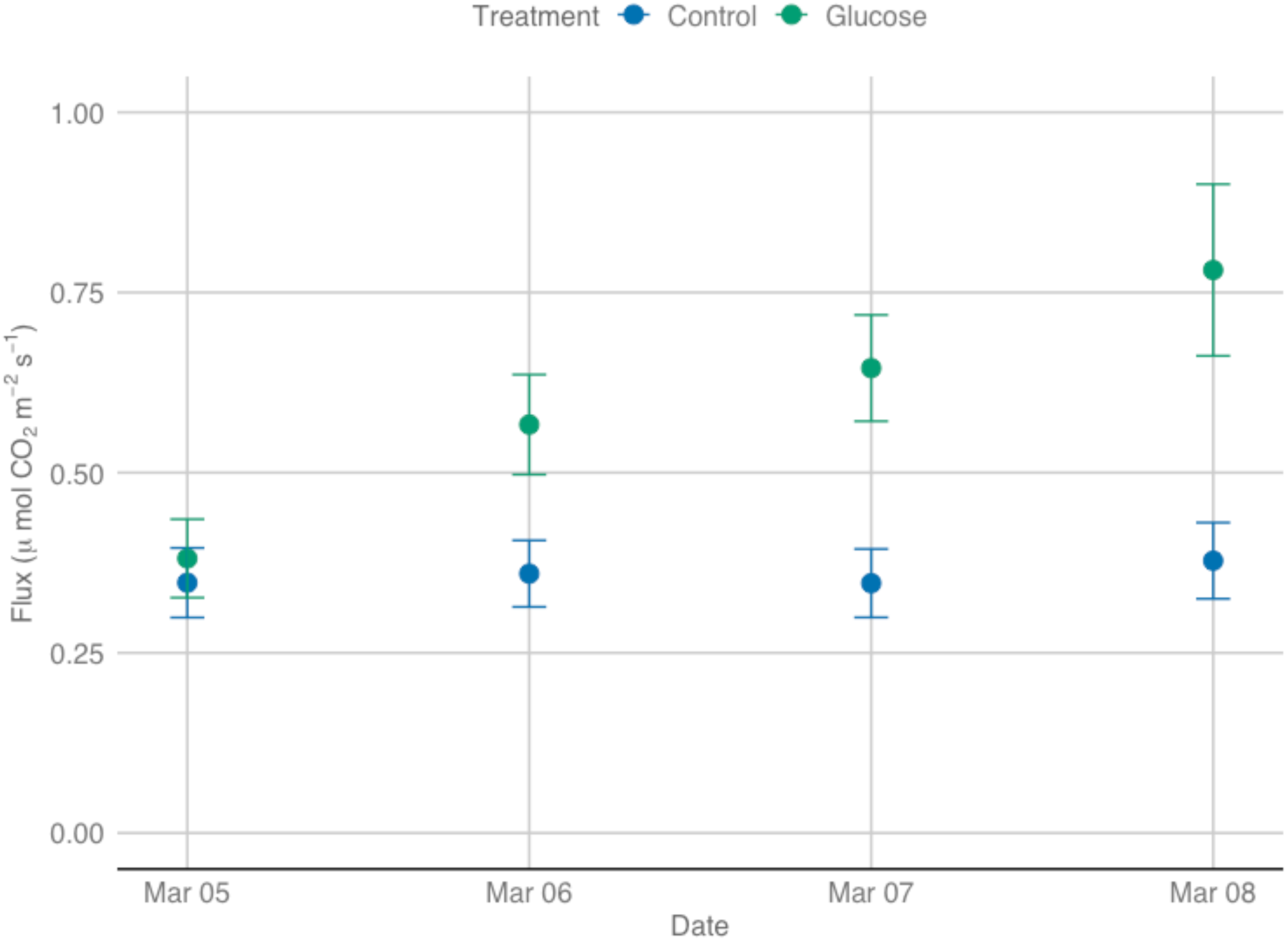
CO_2_ flux response to experimental glucose addition over time in a mixed white spruce, black spruce, paper birch forest in Anchorage, Alaska. Pre-treatment CO_2_ flux measurements were made on March 5, 2019 and were immediately followed by snowpack removal from paired control (n=5) and treatment plots (n=5). Powdered glucose was added to the treatment plots (100 g C/m^2^) and the snowpack was returned to both control and treatment plots. Measurements of CO_2_ flux were made using the diffusion method at 24, 48 and 72 hours after glucose addition. Snow pits were excavated for measurements of snow density and temperature after the final round of measurements and applied to the [CO_2_] measurements made at 24 and 48 hours. Daytime maximum and nighttime minimum air temperatures were consistent over the 72-hour period. Soil surface temperature was −2°C at the beginning and end of the experiment. Bars are S.E.M.

**Fig. S2.**
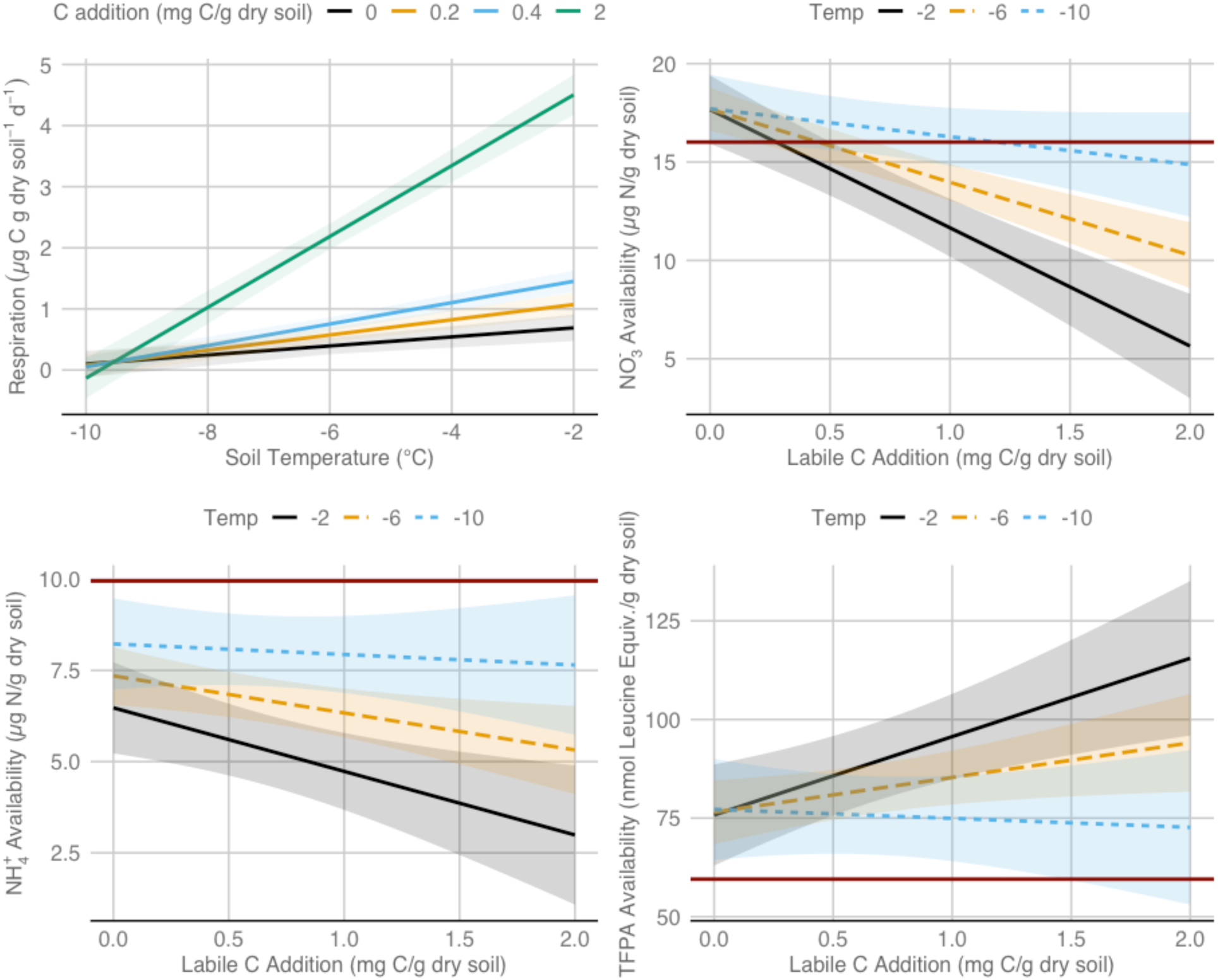
Microbial respiration and soil nutrient availability responses to labile C addition in laboratory incubations at sub-zero soil temperatures. Cellobiose was added at rates of 0, 0.2, 0.4, and 2 mg C per g dry soil to a homogenized composite of root-free organic soil from our hydric and xeric sites and each was separately incubated at −10, −6, −2, 2, and 6 °C (n=4/temperature*C addition treatment). The respiration data were analyzed using a linear mixed effects model, as there was little evidence of non-linearity between −10 and −2°C. Respiration measurements were made approximately weekly during the incubations. Soil nutrient availability was measured at the beginning and end of the three-month incubations. The dark red reference line indicates pre-incubation nutrient availability. Grey shading indicates the S.E.M.

**Fig. S3.**
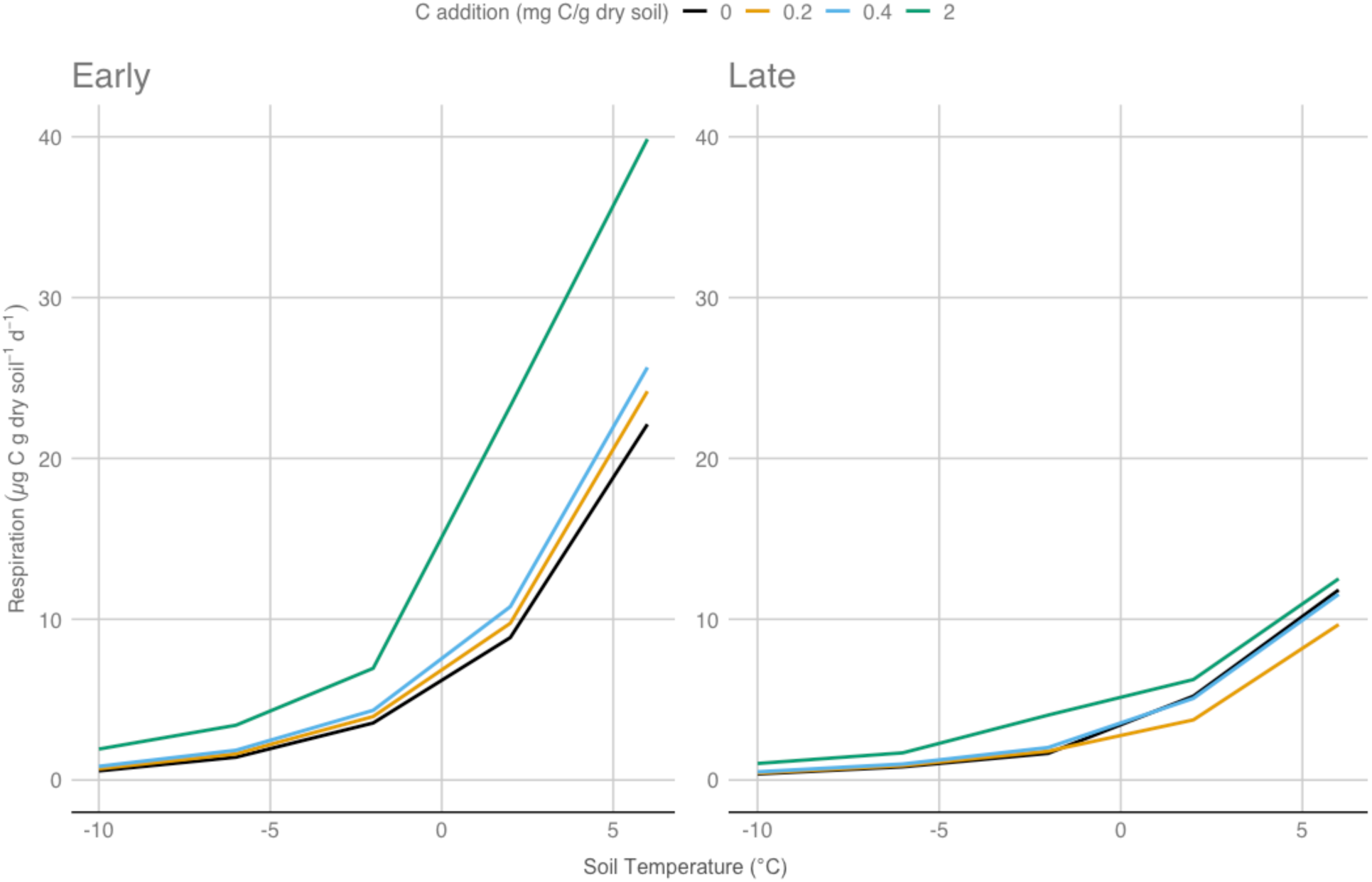
Microbial respiration responses to temperature and labile C addition early and late during the three-month incubation. The incubation dataset was divided approximately in half and a non-linear mixed effects model was fit separately to each half (early versus late). The figure shows model predictions for each combination of temperature and labile C addition level during the two time periods. The interaction between labile C addition and temperature was retained in the model for the late period, in the interest of comparability with the early period, even though there was limited evidence of an interaction.

**Fig. S4.**
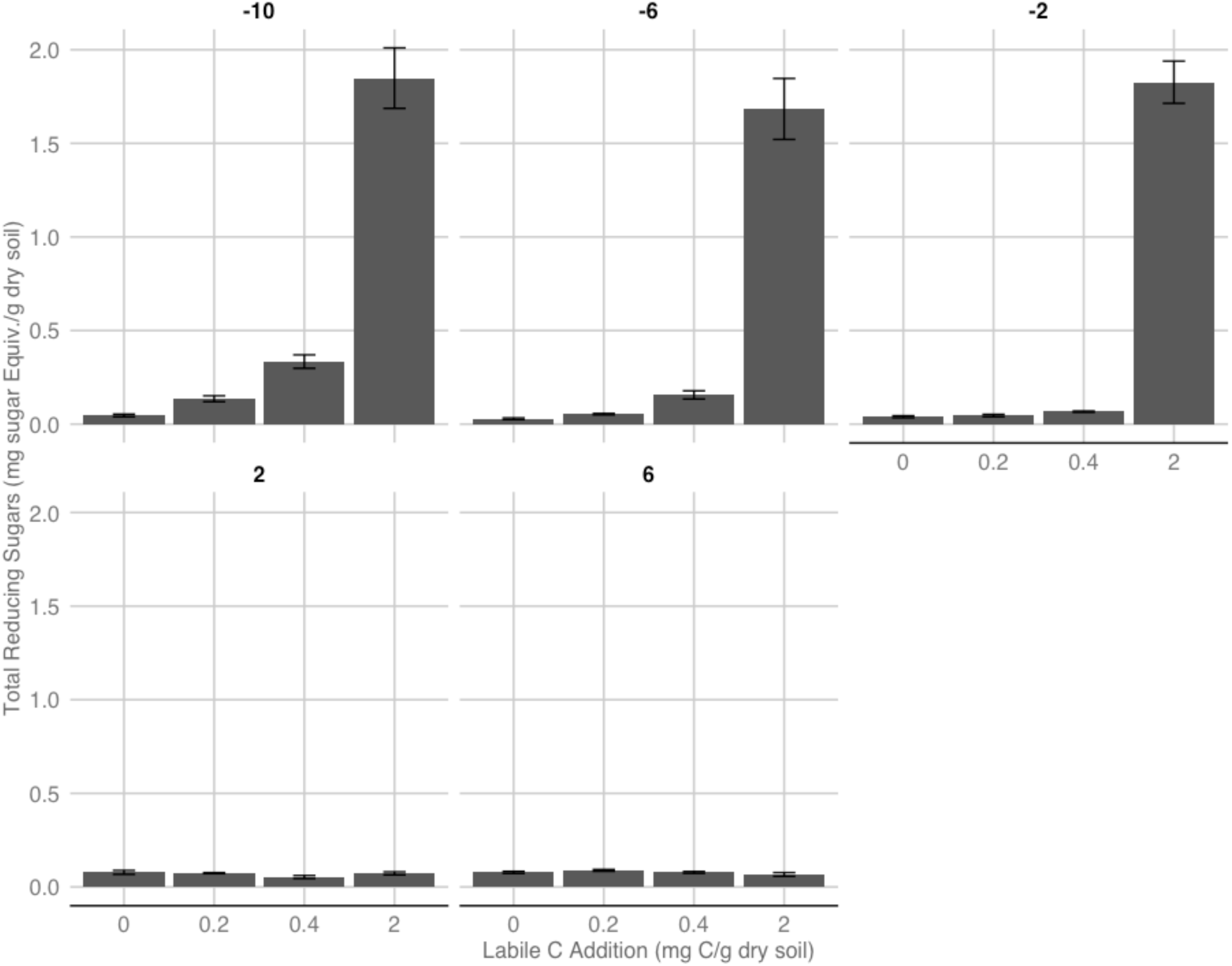
Total reducing sugar (TRS) concentrations in response to variation in temperature and labile C availability. Labile C additions were made using cellobiose, which is a reducing sugar. TRS assays reveal increasing microbial utilization of the added labile C with increasing temperature, including temperature increases below 0°C. Laboratory incubations were performed over three months using homogenized composite organic soils collected from our hydric and xeric sites (n=4/temperature*C addition treatment). Bars are 1.0 S.E.M.

**Fig. S5.**
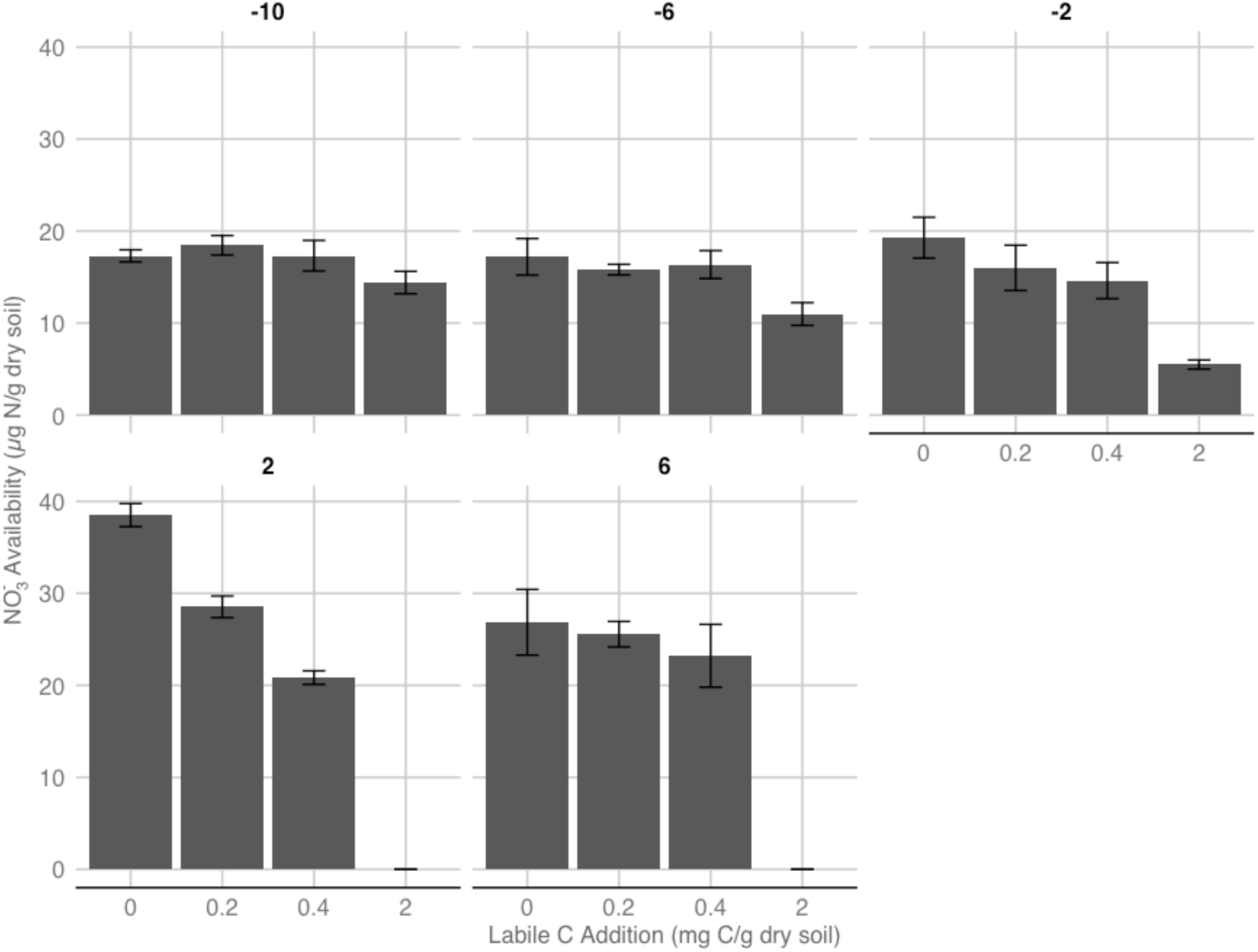
NO_3_^-^ availability in response to variation in temperature and labile C availability. Laboratory incubations were performed over three months using homogenized composite organic soils collected from our hydric and xeric sites (n=4/temperature*C addition treatment). Bars are 1.0 S.E.M.

**Fig. S6.**
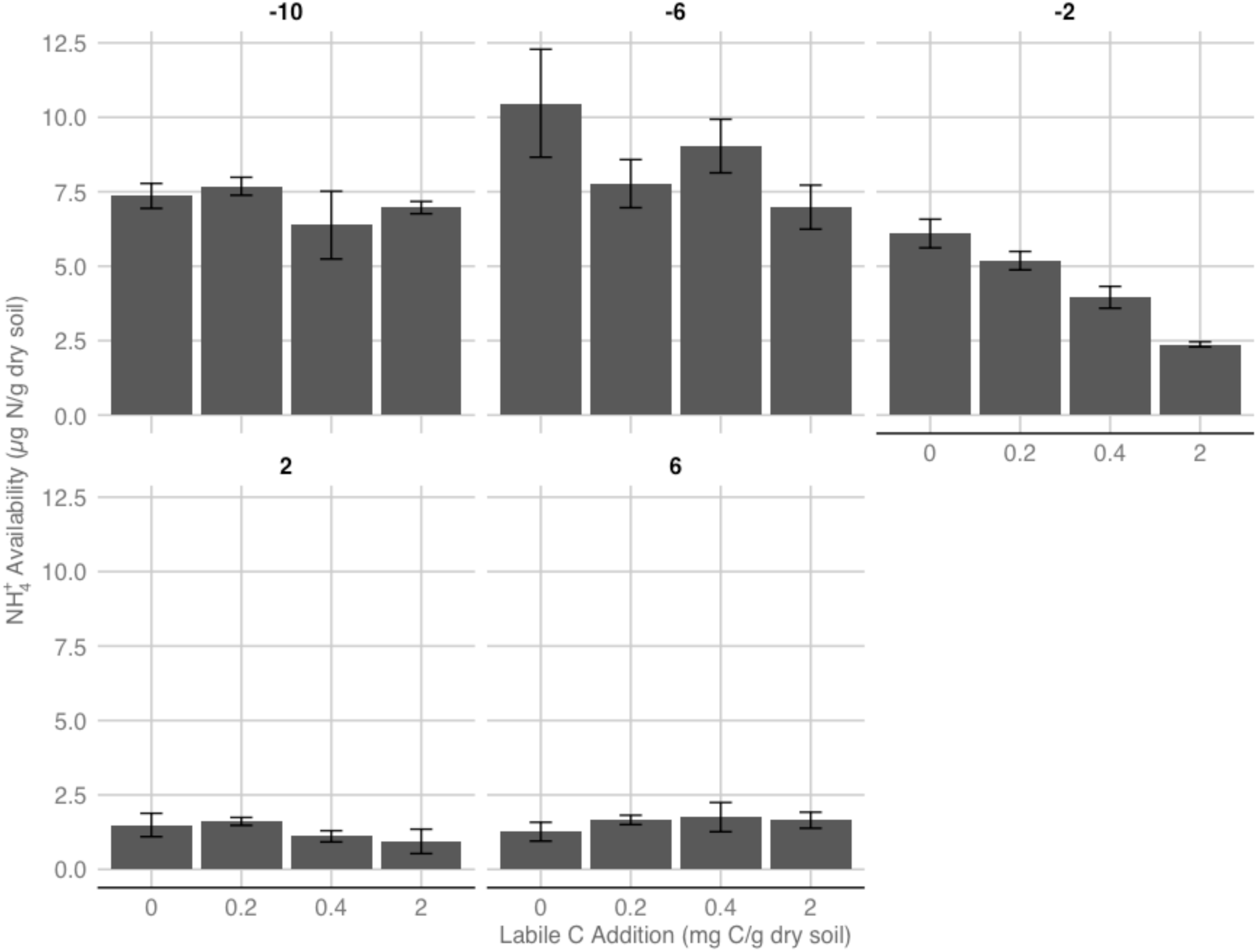
NH_4_^+^ availability in response to variation in temperature and labile C availability. Laboratory incubations were performed over three months using homogenized composite organic soils collected from our hydric and xeric sites (n=4/temperature*C addition treatment). Bars are 1.0 S.E.M.

**Fig. S7.**
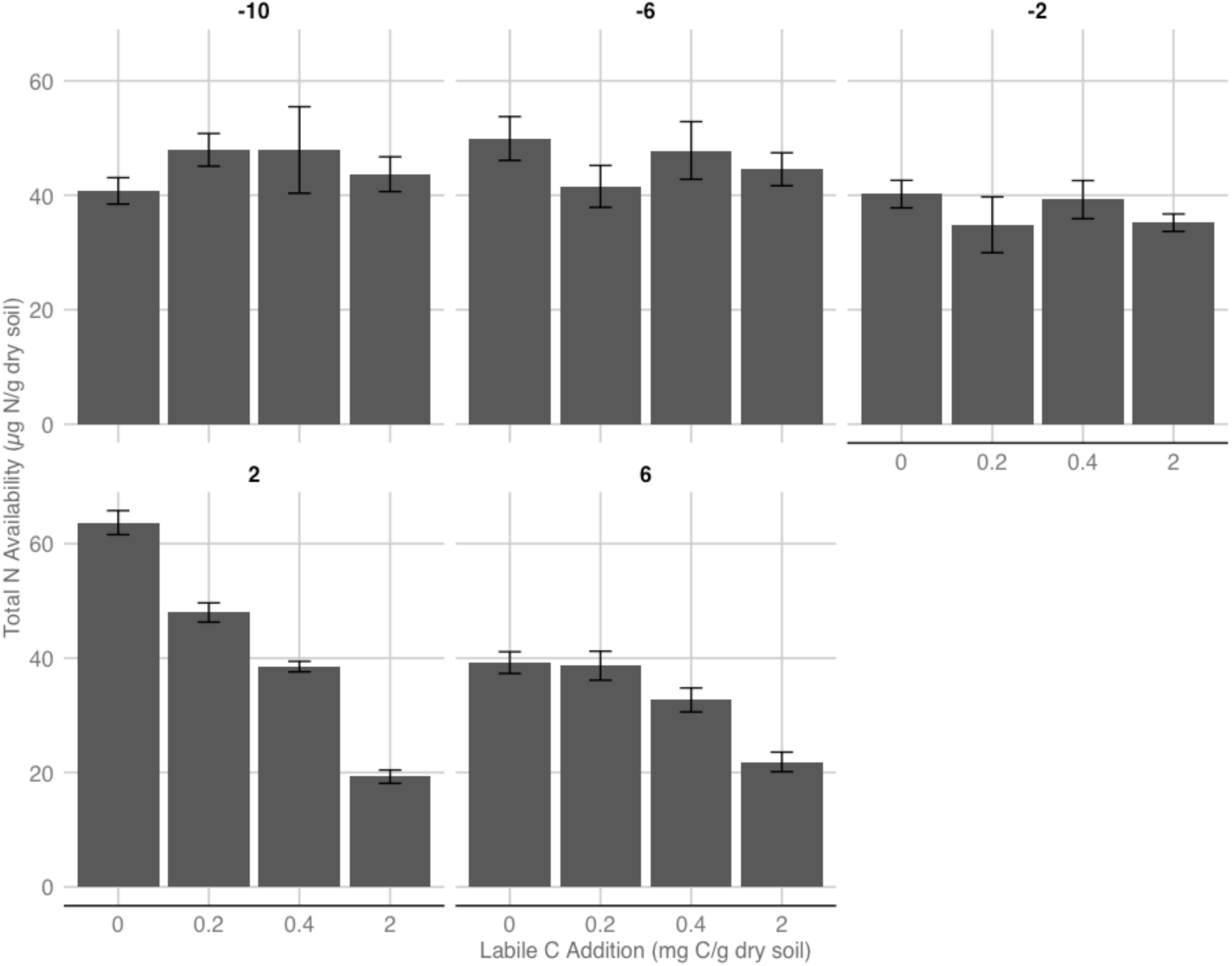
Total nitrogen (N) availability in response to variation in temperature and labile C availability. Laboratory incubations were performed over three months using homogenized composite organic soils collected from our hydric and xeric sites (n=4/temperature*C addition treatment). Bars are 1.0 S.E.M.

**Fig. S8.**
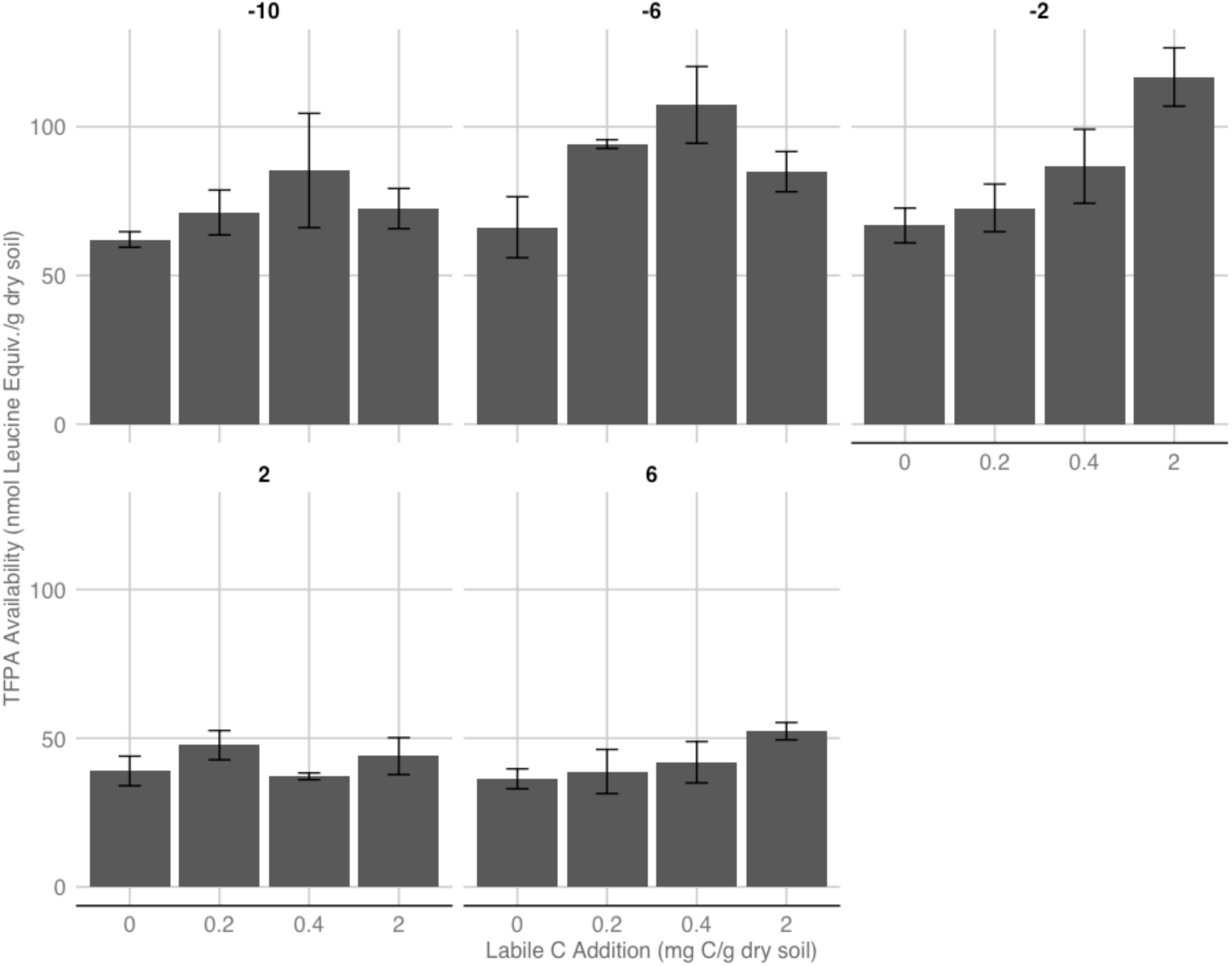
Total free primary amine (TFPA) availability in response to variation in temperature and labile C availability. TFPA is an indicator of amino acid availability. Laboratory incubations were performed over three months using homogenized composite organic soils collected from our hydric and xeric sites (n=4/temperature*C addition treatment). Bars are 1.0 S.E.M.

**Fig. S9.**
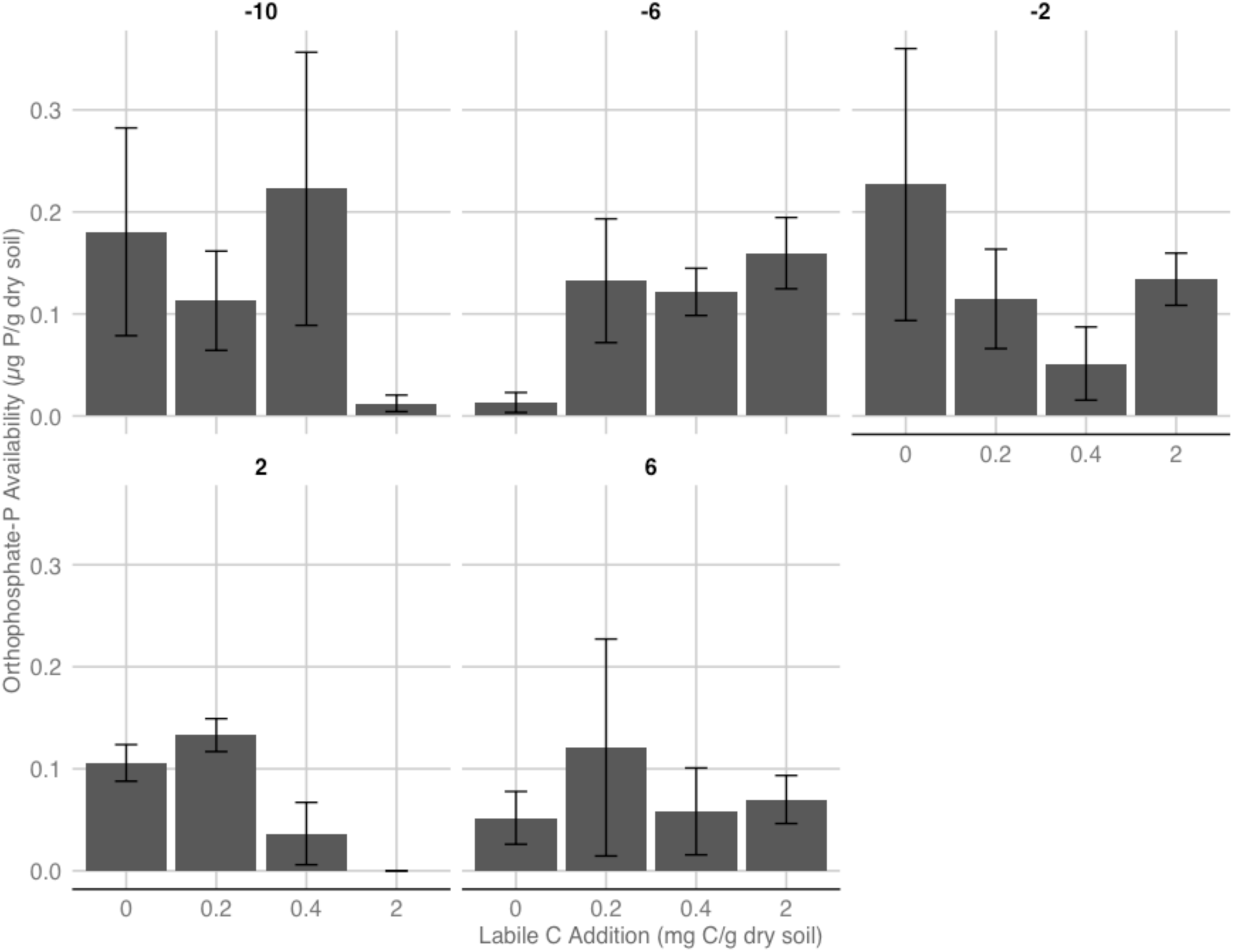
Orthophosphate-P availability in response to variation in temperature and labile C availability. Laboratory incubations were performed over three months using homogenized composite organic soils collected from our hydric and xeric sites (n=4/temperature*C addition treatment). Bars are 1.0 S.E.M.

**Fig. S10.**
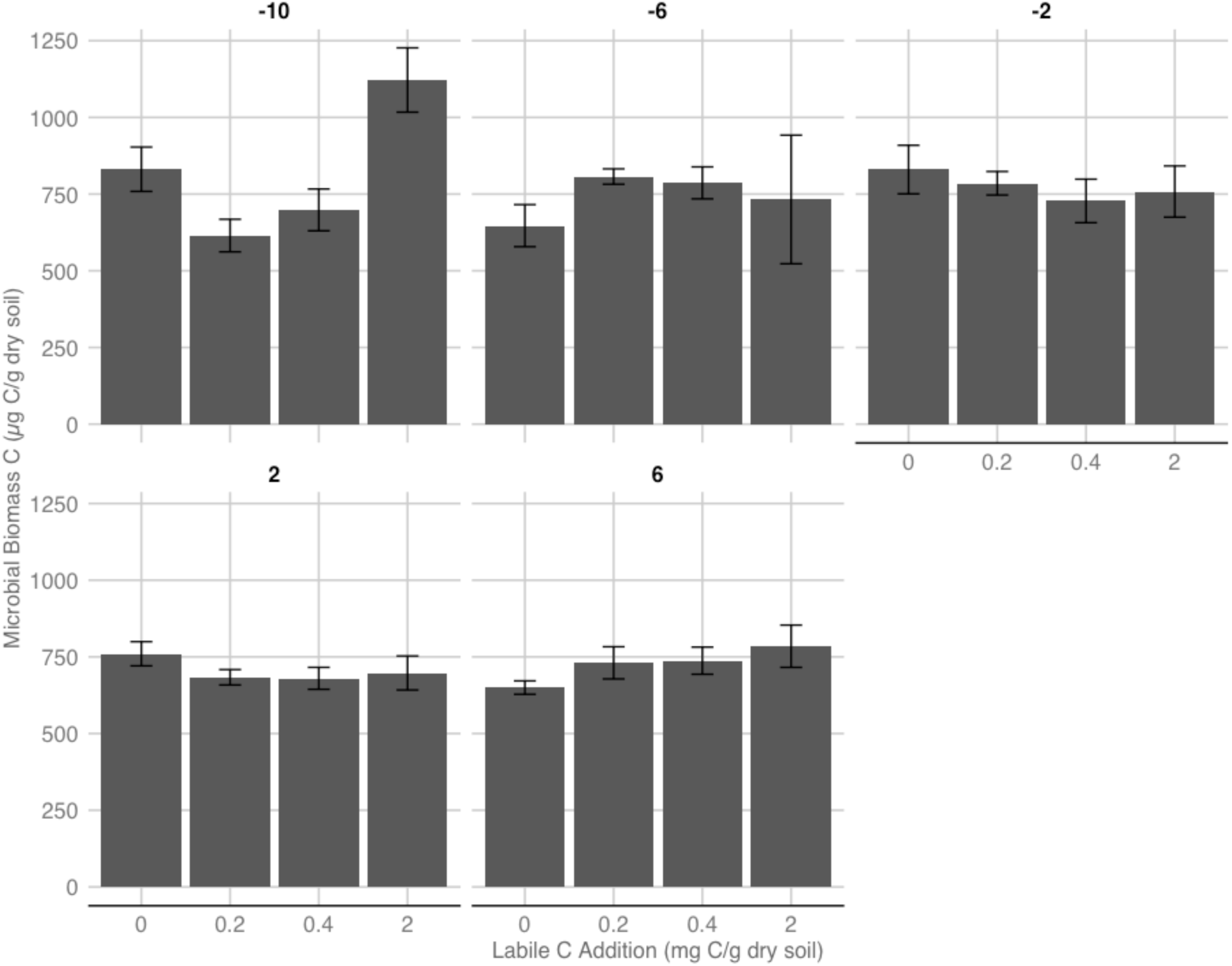
Microbial biomass carbon (C) in response to variation in temperature and labile C availability. Laboratory incubations were performed over three months using homogenized composite organic soils collected from our hydric and xeric sites (n=4/temperature*C addition treatment). Bars are 1.0 S.E.M.

**Fig. S11.**
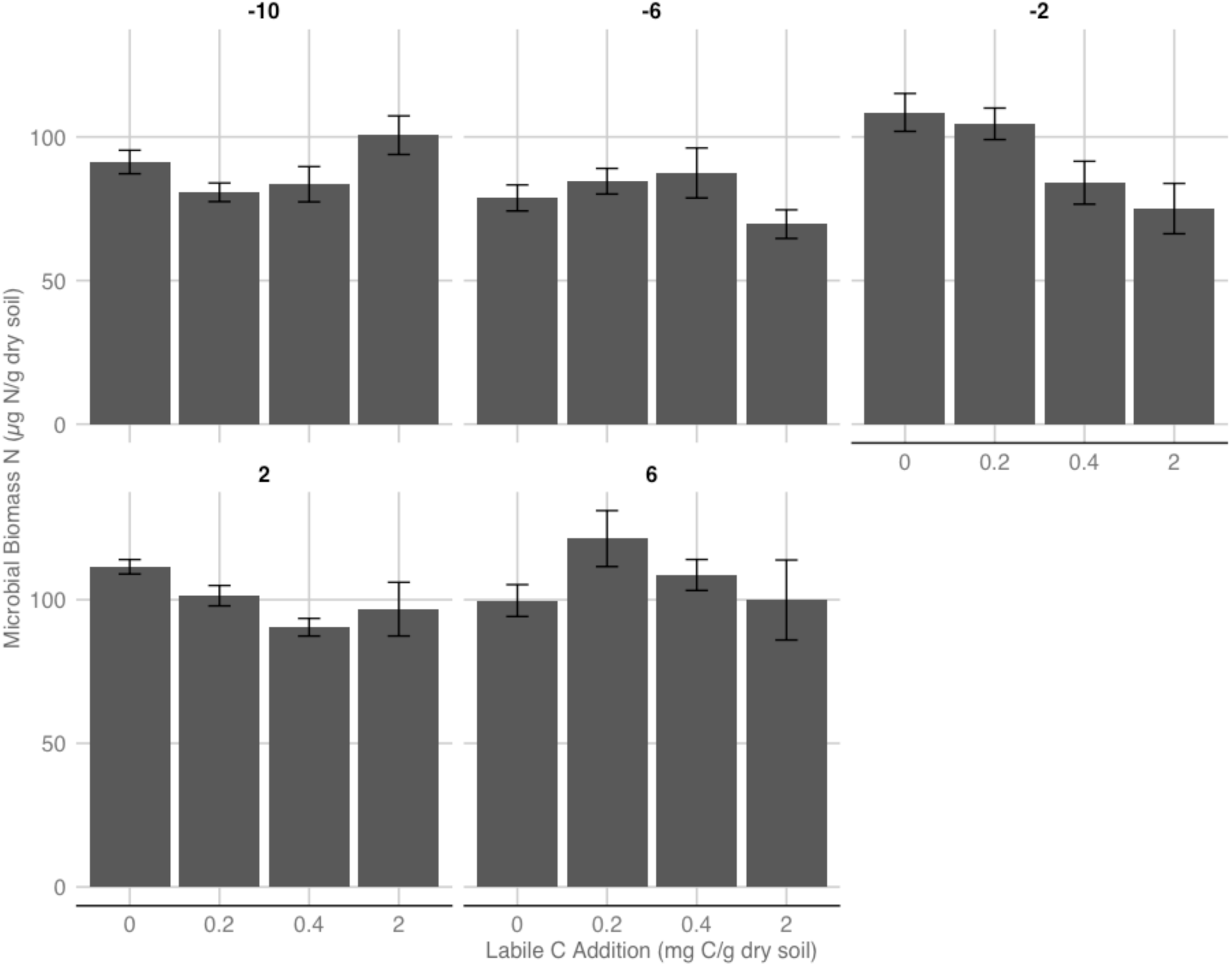
Microbial biomass nitrogen (N) in response to variation in temperature and labile C availability. Laboratory incubations were performed over three months using homogenized composite organic soils collected from our hydric and xeric sites (n=4/temperature*C addition treatment). Bars are 1.0 S.E.M.

